# Small Vessel Diseases: 3D Characteristics of the Vasculature and White Matter

**DOI:** 10.1101/2021.06.30.450101

**Authors:** Rie Saito, Kazuki Tainaka, Hiroaki Nozaki, Masahiro Uemura, Yasuko Toyoshima, Masahiro Suzuki, Masaharu Tanaka, Arika Hasegawa, Takashi Abe, Aki Sato, Hideki Hashidate, Shuichi Igarashi, Ryoko Koike, Akihiko Ueda, Mitsuharu Ueda, Yukio Ando, Kohei Akazawa, Osamu Onodera, Akiyoshi Kakita

## Abstract

Cerebral small vessel disease (SVD) is associated with white matter hyperintensities (WMHs), thereby contributing to vascular dementia and movement disorder. However, the pathomechanisms responsible for WMH-related small vessel degeneration remain poorly understood due to the technical limitations of current methods. The aim of this study was to clarify the 2- and 3-dimensional (2D and 3D) pathological features of small vessels and white matter (WM) in the brains of patients with cerebral autosomal dominant arteriopathy with subcortical infarcts and leukoencephalopathy (CADASIL), *HTRA1*-autosomal dominant disease (*HTRA1*-AD) and sporadic SVD (sSVD). From a cohort of 86 consecutive autopsied patients with SVDs, we retrieved those with genetically confirmed CADASIL and *HTRA1*-AD (three and three, respectively), and four with sSVD. We quantitatively evaluated WM and vascular changes in the frontal portion of the centrum semiovale and temporal lobe using conventional 2D and chemically cleared 3D analytical methods with light-sheet fluorescence microscopy. Quantitatively, the WM pathology, including the density of myelin, axons and gliosis, was most severe in CADASIL, but unexpectedly sSVD was second in order of severity, followed by *HTRA1*-AD. The density of clasmatodendrocytes, known to be irreversibly injured astrocytes, was considerably highest in *HTRA1*-AD. The vascular pathology, including arteriole and capillary sclerosis and the extent of the perivascular space, was most severe in CADASIL, whereas the density of smooth muscle actin (SMA) positivity was most decreased in *HTRA1*-AD. 3D immunohistochemistry for SMA demonstrated two distinct patterns of SMA loss within the vessels: (1) CADASIL and sSVD: diffuse loss, being prominent in small branches, (2) *HTRA1*-AD: selective loss in main branches. Overall, the extent of WM and vascular degeneration is most severe in CADASIL, whereas SMA loss is most evident in *HTRA1*-AD. These differences in the size and distribution of affected vessels may be related to the heterogeneous WM pathology and underlying pathomechanisms of SVD.

## Introduction

Cerebral small vessel disease (SVD) is a sporadic or hereditary condition affecting cerebral arterioles, capillaries and venules, being a major cause of vascular cognitive impairment (VCI) characterized by multiple lacunar infarcts, leukoaraiosis and gait disturbance.^1–5^ These neuronal and movement disorders are associated with white matter hyperintensities (WMHs) evident on T2-weighted MRI,^5,6^ suggesting that the pathomechanisms of small vessel degeneration act on white matter (WM) networks. However, the pathological basis of WMHs remains poorly understood.^7,8^ *In vivo*, it is still challenging to differentiate SVD-related pathology from coexisting conditions such as Alzheimer’s disease,^8^ and to visualize small vessels. Moreover, imaging of the actual three-dimensional (3D) vasculature in small thin-tissue sections is impractical. Therefore, structural alterations of small vessels have remained unclear, even in the sporadic form, which affects most patients with SVD.

In contrast, recent genetic studies have provided molecular information useful for better understanding both the sporadic and hereditary forms. Cerebral autosomal dominant arteriopathy with subcortical infarcts and leukoencephalopathy (CADASIL) is the most common heritable cause of stroke and vascular dementia in adults caused by mutations in the *NOTCH3* gene.^9,10^ The clinicopathological features of patients with CADASIL resemble those in sporadic SVD,^7,10–12^ except for specific abnormal deposition of the extracellular domain of Notch3 within vessel walls.^13^ Recently, in stroke survivors and CADASIL patients, it has been proposed that clasmatodendrocytes, known to be irreversibly injured astrocytes, could be a marker associated with WMH and gliovascular interactions at the blood-brain barrier.^4,15^ However, little is known about interactions between clasmatodendrocytes and changes in small vessels and the WM.

Cerebral autosomal recessive arteriopathy with subcortical infarcts and leukoencephalopathy (CARASIL) is an autosomal recessive inherited cerebral SVD characterized by leukoaraiosis, alopecia and spondylosis deformans caused by mutations in the High temperature requirement serine peptidase A1 (*HTRA1*) gene.^16^ CARASIL appears to be much less prevalent than CADASIL, whereas heterozygous mutations in *HTRA1* have been identified recently in some families showing SVD (*HTRA1*-autosomal dominant disease, *HTRA1*-AD),^17,18^ suggesting that these are more frequent and important than previously considered. As compared with CARASIL, similar but milder features of vasculopathy such as a multi-laminated internal elastic lamina with severe loss of smooth muscle actin (SMA) have been reported in a few patients,^19–21^ but the detailed pathological features of *HTRA1*-AD remain largely elusive.

In the present study, using a 2D quantitative method, we analyzed the pathological features associated with integrity of the WM and vascular networks in sporadic pure SVD, CADASIL and *HTRA1*-AD to clarify their relationship with small vessel and WM degeneration. Conventional 3D reconstitution of thinly sliced 2D serial sections facilitates cellular-level resolution, but is labor-intensive and not practical. To further examine the spatial distribution of SMA loss within vascular networks and gliosis in the degenerative WM, we improved a novel 3D analytical method reported previously using chemical tissue clearing protocols and light-sheet fluorescence microscopy, allowing 3D imaging of large immunostained specimens of human brain tissue including the WM.^22,23^

## Materials and Methods

### Subjects

We reviewed the medical records and neuropathologic features of 86 consecutive patients with pathologically proven severe cerebral arteriolosclerosis (48 males and 38 females; aged 73.8±11 [mean±SD; range 46-95 years]) who were referred to the Brain Research Institute, Niigata University, between 1965 and 2018. This cohort did not include any patients with ischemic or hypoxic brain. We selected 82 patients for whom sections of all 4 cerebral lobes and basal ganglia were available. To identify the patients with CADASIL and *HTRA1*-AD, we pathologically examined all patients to check whether there were specific histopathologic findings including abnormal deposition of the extracellular domain of Notch3 in CADASIL or multi-laminated internal elastic lamina in CARASIL, and performed genetic analysis of the 40 patients whose samples were available for the test. As a result, we retrieved three patients with cerebral autosomal dominant arteriopathy with subcortical infarcts and leukoencephalopathy (CADASIL group), and three with *HTRA1*-autosomal dominant disease (*HTRA1*-AD group) (Table 1). The present series of patients did not include any with CARASIL. The detailed clinicopathologic features of *HTRA1*-AD 1 have been described previously.^21^ Among patients with sporadic pure SVD, we excluded 31 showing cognitive decline or movement disorder due to comorbid pathologies (Alzheimer’s disease = 11, Parkinson’s disease = 8, progressive supranuclear palsy = 3, cerebral amyloid angiopathy = 2, amyotrophic lateral sclerosis = 2, myotonic dystrophy = 2, severe argyrophilic grain disease = 1, myopathy = 1 and HTLV-1 associated myelopathy = 1), ^25^ leaving a cohort of 45 patients with cerebral small vessel disease (CSVD) type 1 (age-related and vascular risk factor-related).^1^ Of these patients, we identified 13 with subtype II (multiple small or lacunae, small vessel disease) based on the Newcastle categorization.^8,24^ The other patients were classified as follows: subtype I (large infarcts or cortical infarcts) = 12, subtype III (strategic infarcts) = 10, subtype IV (hypoperfusive lesions, hippocampus sclerosis) = 0, and subtype V (cerebral hemorrhage) = 10.^8,24^ We then included four of the 13 patients considering the sufficiency of the clinical and genetic information (Table 1). Because the patients with CADASIL and *HTRA1*-AD were apparently younger than those with sSVD (58.7±12.3, 65.0±3.6 and 81.3±3.5), we also included individuals who were age-matched to the patients with hereditary SVD without any neurological disorders as a control group (59.7±7.4, n = 3, Con 1-3 in Table 2B). The present study was approved by the institutional review board of Niigata University (G2018-0010), Niigata, Japan. Informed consent for autopsy including the use of tissues for research purposes was obtained from the patients’ families.

**Table 1.**
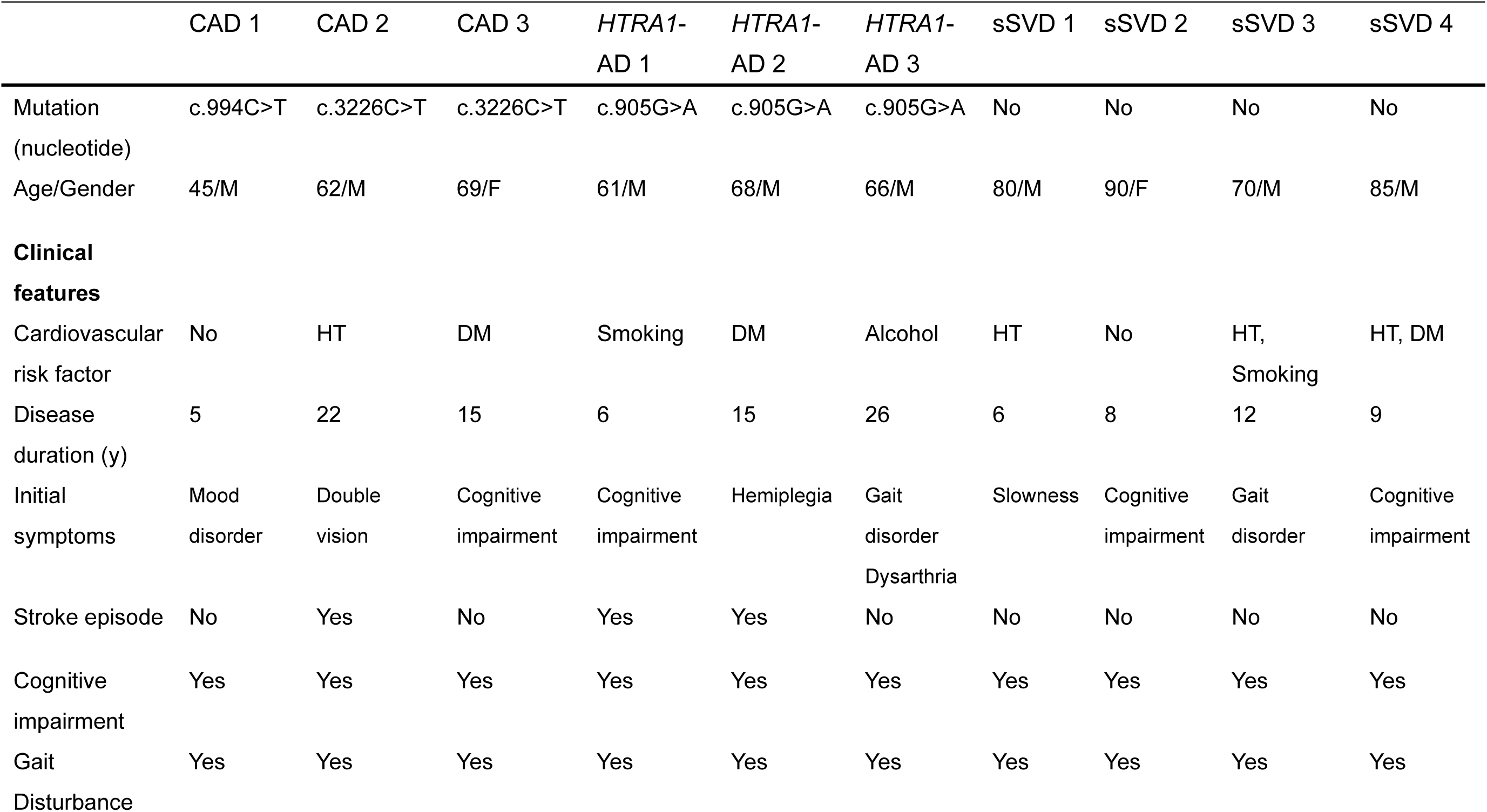
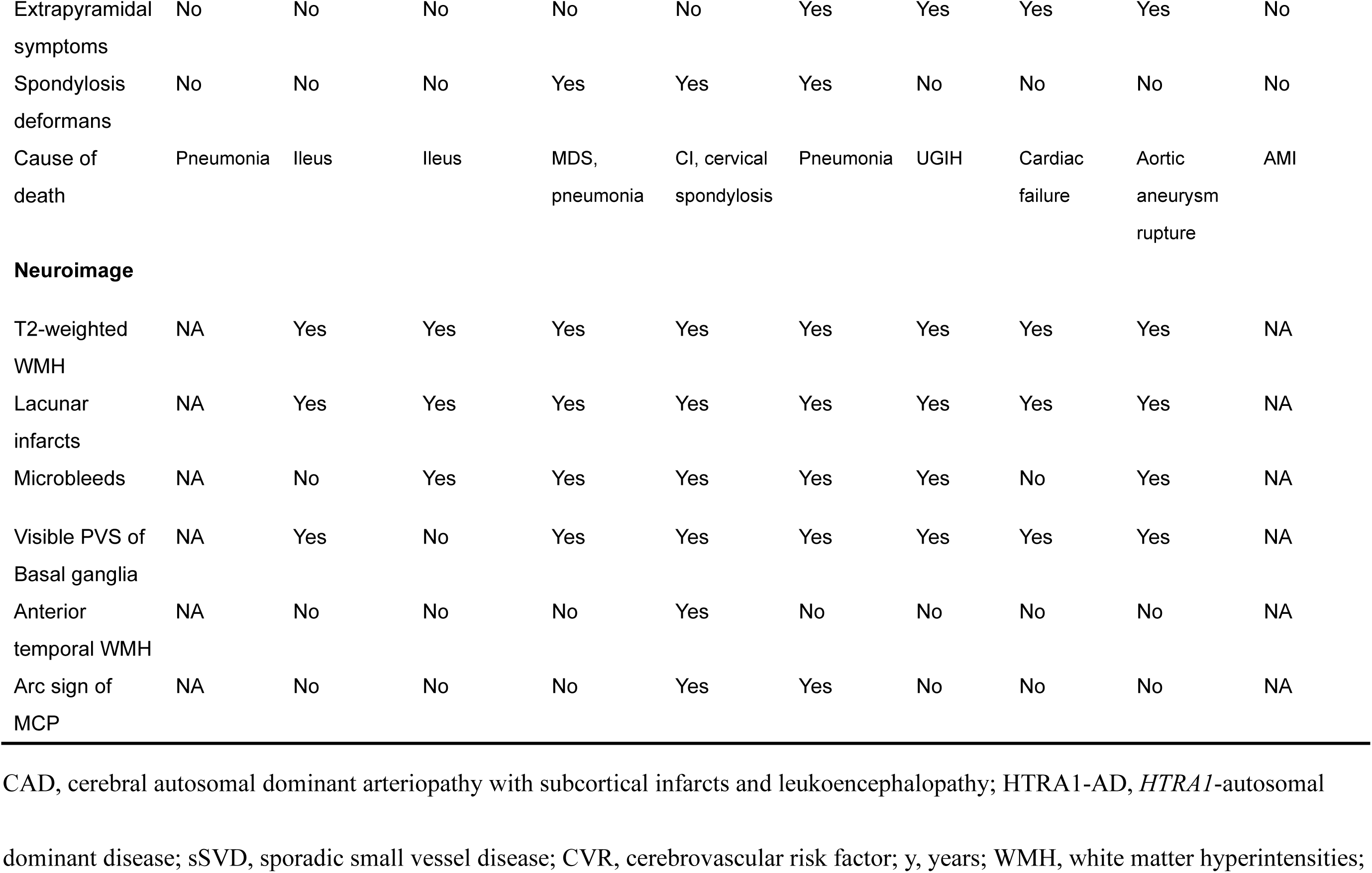
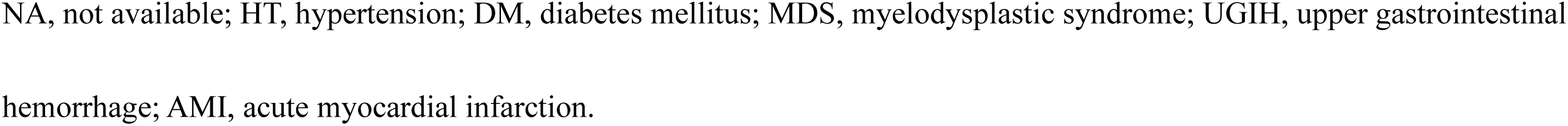
Clinical and neuroimaging features in the patients with cerebral small vessel diseases.

**Table 2.**
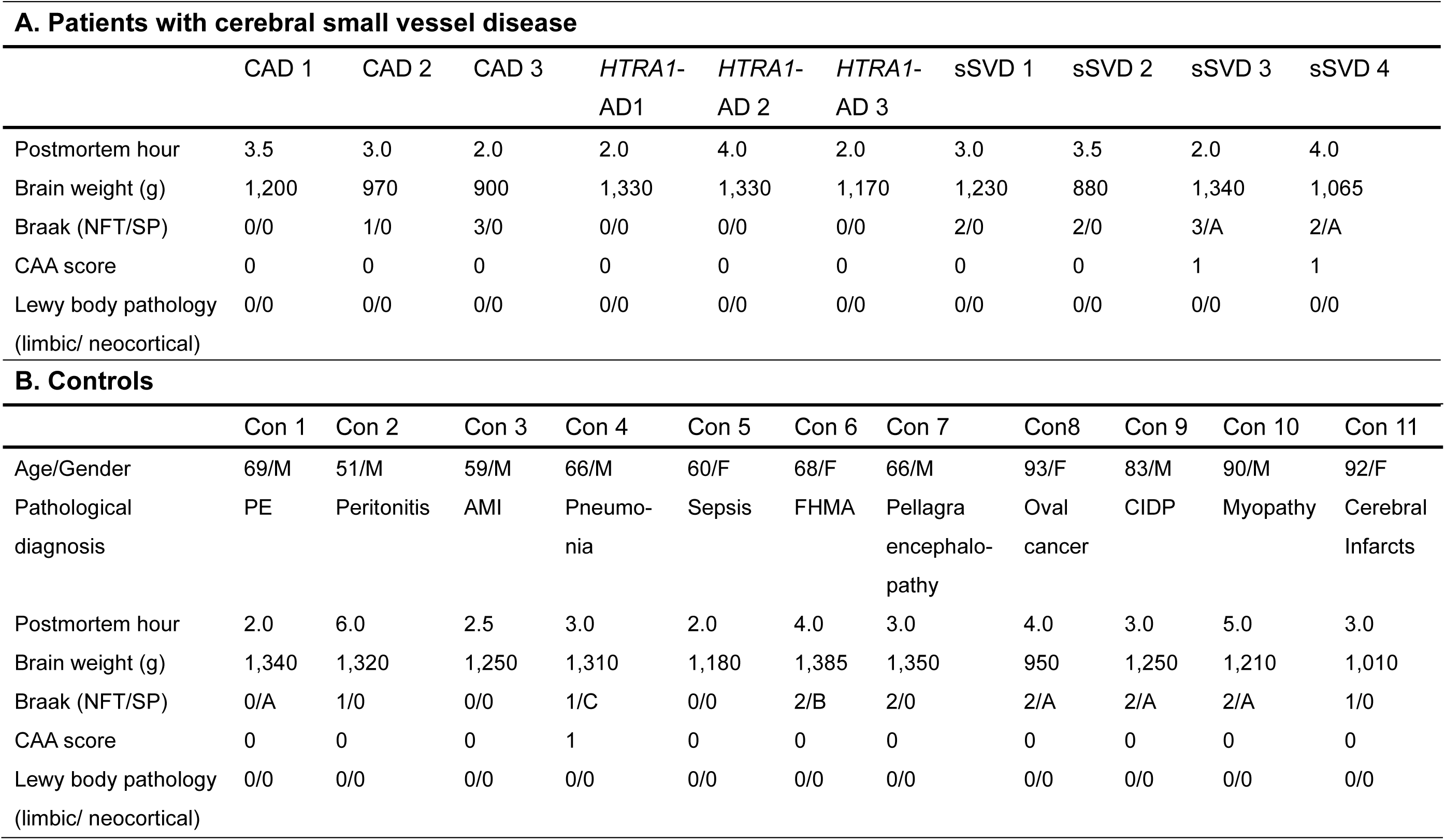
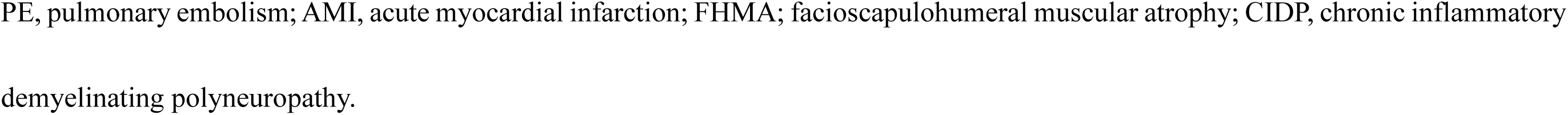
Neuropathological profile in the patients with cerebral small vessel diseases and controls

### Patients’ Profile

The clinical and pathological profiles of the patients are summarized in Table 1 and Table 2A.

Genetic analysis detected two heterozygous missense mutations – c.994C>T and c.3226C>T in the *NOTCH3* gene – and one heterozygous missense mutation – c.905G>A – in the *HTRA1* gene in the patients with CADASIL and *HTRA1*-AD, respectively. All mutations were reported in patients with CADASIL and *HTRA1*-AD.^18,26,27^

Two of the three patients with CADASIL, all three with *HTRA1*-AD, and three of the four with sSVD had cerebrovascular risk factors such as hypertension, diabetes mellitus, a history of smoking or a history of heavy alcohol consumption. The age at onset and disease duration in the CADASIL and *HTRA1*-AD groups were similar (45.0 ±7.8 and 49.3±8.1 years, and 13.7±9.1, 15.7±8.2 years), whereas those in the sSVD group were much later and shorter (72.5±10.3 years and 8.8±2.5 years). Cognitive impairment and gait disturbance were present in all patients in the three groups. Interestingly, one of the three patients with *HTRA1*-AD and three of the four patients with sSVD presented with extrapyramidal symptoms such as tremor or slowness despite absence of Lewy body disease or other Parkinson-related neurodegenerative diseases (Table 2). All of the patients with *HTRA1*-AD suffered from severe spondylosis deformans, known to be a hallmark of CARASIL or *HTRA1*-AD, and one of these patients died of cervical spondylosis.

Brain MRI revealed leukoaraiosis with multiple lacunar infarcts in the deep WM in all ten patients for whom data were available. Based on Fazekas’ rating scale, both the periventricular hyperintensity and separate deep WM hyperintense signals in all the patients were rated as 3 (irregular PVH extending into the deep white matter and large confluent areas, respectively), i.e. the most severe score^28^ (Fig 2A-C). Almost all patients had microbleeds, and dilatation of the PVS detected as dot or linear structures with T2-weighted hyperintensity in the basal ganglia (Fig 2D-E, F). The number of visible PVSs was high in the patients with *HTRA1*-AD and sSVD, and moderate in those with CADASIL. Anterior temporal hyperintensities were detected in one patient with *HTRA1*-AD (Fig 2E, *upper right panel*), and arc-shaped hyperintensities in the middle cerebellar peduncles were detected in two of the three patients with *HTRA1*-AD (Fig 2E, *lower right panel*), as observed commonly in CARASIL.^29^

Neuropathological analysis demonstrated that no patients with CADASIL or *HTRA1*-AD had age-associated pathology such as Alzheimer’s disease, CAA or Lewy body pathology (Table 2A).

### Neuropathological Analysis and Immunohistochemistry

All of the brains were fixed in formalin and processed as described previously.^30^ All patients were assessed for structural integrity and infarcts and myelin loss using 4-μm-thick sections stained with hematoxylin and eosin and Klüver-Barrera (KB), respectively. Macroscopically, WM atrophy was evident predominantly in the frontal CSO in the CADASIL, *HTRA1*-AD and sSVD patients with longer disease duration, while the cortices were relatively preserved. Consistent with the MRI finding of severe leukoencephalopathy in these groups, myelin pallor detected by KB staining was commonly observed in the frontal CSO (Fig 1A), especially the deep WM and periventricular area, where aberrant axons and multiple small infarcts with gliosis were observed, whereas the temporal WM was relatively well preserved. Similarly, we found that arteriolosclerosis was accentuated in the periventricular area of the frontal CSO and basal ganglia, accompanied by capillary change involving loss of endothelial cells with wall thickening in both the hereditary and sporadic SVD groups, although characteristic vascular changes in CADASIL and *HTRA1*-AD such as deposition of basophilic granules with positive immunoreactivity for NOTCH3 ECD in the SMC or multilayering of the internal elastic lamina, extended throughout the brain. Therefore, for quantitative analyses in the present study, we prepared coronal slices of the frontal lobe containing the CSO and temporal lobe at the level of the anterior hippocampus from all the subjects. The slices were embedded in paraffin, and serial 4-μm-thick sections were cut.

**Figure 1:**
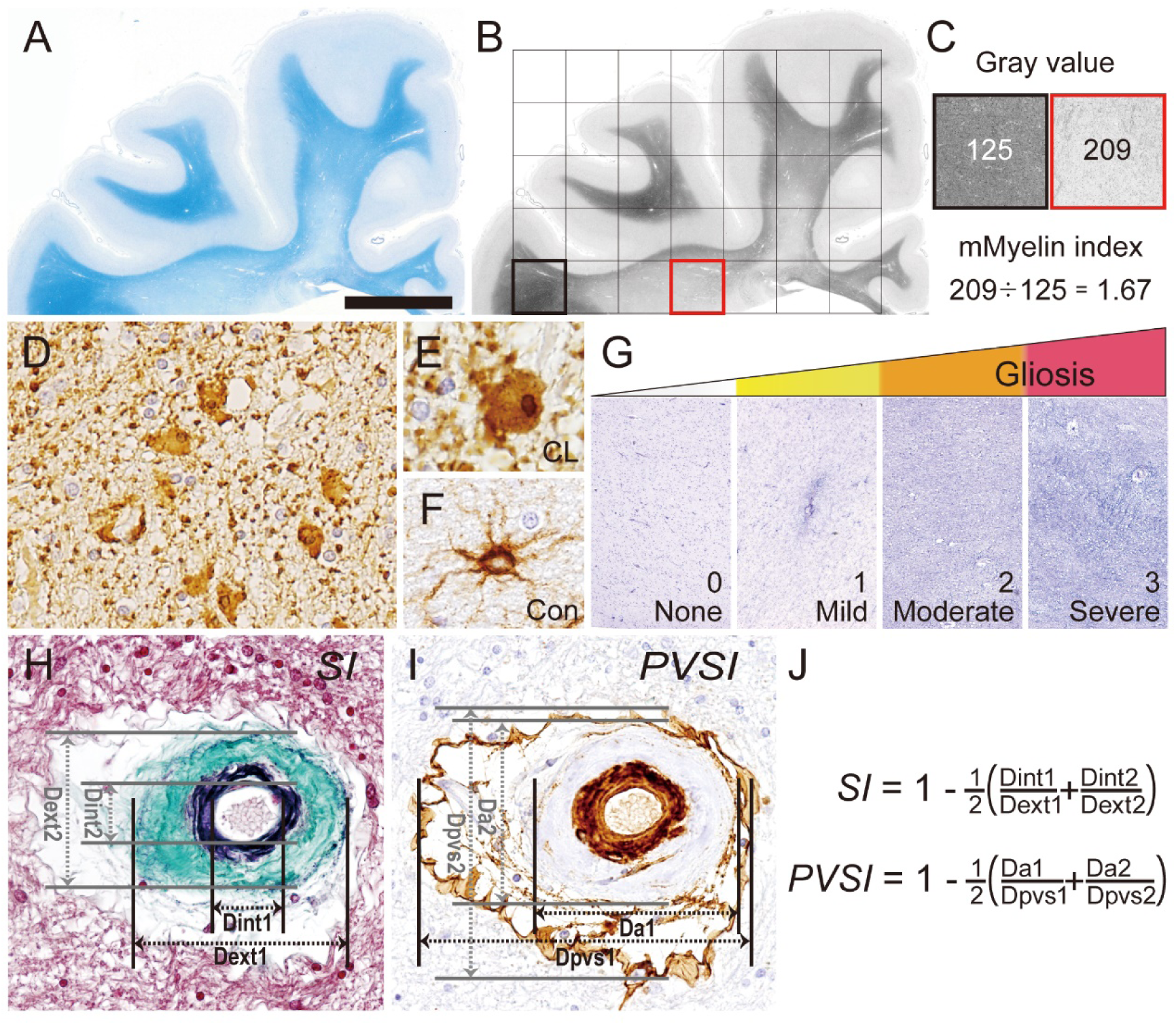
Quantification of the white matter degeneration and vasculopathy. **(A-C)** Calculation of the modified Myelin index (mMyelin index). A captured image of a section subjected to Klüver-Barrera staining **(A)** is converted to a gray scale image with grid squares of 5 × 5 mm^2^ **(B)**. The mMyelin index is the gray value ratio of the apparently normal area (*black*) to the region of interest (r*ed*) **(C)**. **(D-F)** The appearance of GFAP-positive clasmatodendrocytes. Image of a cluster of clasmatodendrocytes **(D)**, and magnified images showing a clasmatodendrocyte **(E)** and an apparently normal astrocyte **(F)**. **(G)** Quartile analysis of gliosis using Holzer-stained sections. **(H-I)** Measurement of arteriole sclerosis and PVS. The lengths of the internal diameter (Dint) and the external diameter (Dext) of an arteriole in cross-section were used to calculate the sclerosis index **(H)**. The lengths of the external diameter of an arteriole (Da) and the external diameter of a PVS were used to obtain the PVS extension index **(I)**. **(J)** Formulae for the sclerosis index and the PVS extension index. CL, clasmatodendrocytes; Con, control; SI, sclerosis index; PVSI, perivascular extension index. A-I, Frontal centrum semiovale. Bar = 1 cm **(A, B)**, 60 μm **(D)**, 40 μm **(E, F)**, 200 μm **(G)** and 10 μm **(H, I)**

**Figure 2:**
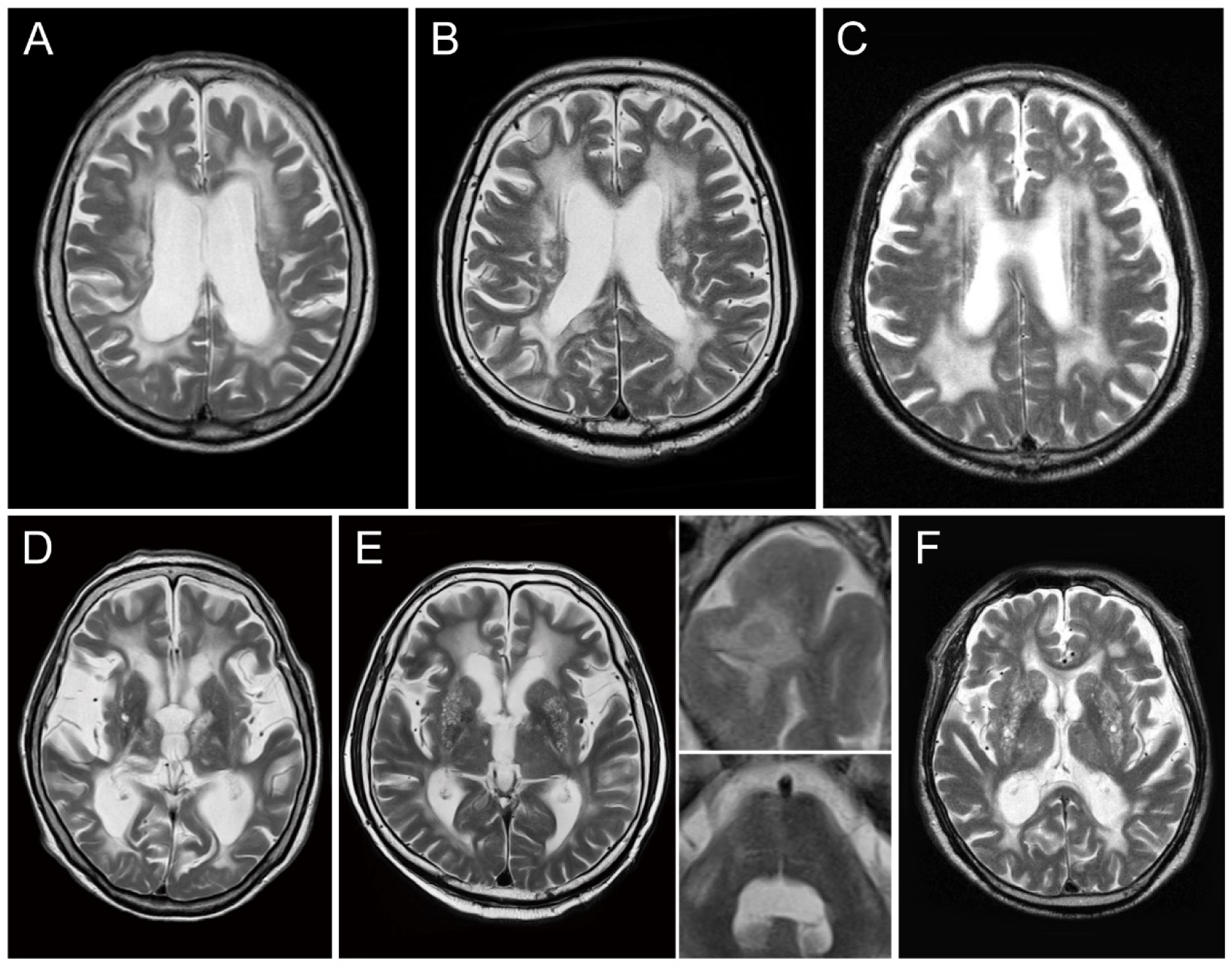
Representative brain MRI findings of the patients with small vessel diseases. Axial view of T2-weighted images. **(A, D)** A patient with CADASIL (CAD 3 in Table 1). **(B, E)** A patient with *HTRA1*-AD (*HTRA1*-AD 2). **(C, F)** A patient with sporadic SVD (sSVD 1). **(A-C)** Confluent white matter hyperintensities extend from the periventricular area to subcortical white matter, predominantly in the frontal centrum semiovale. “Central atrophy” showing predominant atrophy of the central white matter with lateral ventricle dilation. **(D-F)** The number of punctate or linear structures in the basal ganglia regarded as enlarged PVS varies among the patients. In the patient with CADASIL, bilateral linear hyperintensities are evident in the external capsules **(D)**. Hyperintensities in the anterior temporal lobe **(E, *upper right panel*)** and an arc-shaped lesion in the middle cerebellar peduncles **(E, *lower right panel*)**.

The sections of the frontal CSO and temporal lobe were stained using several methods: Klüver-Barrera, Holzer, and elastica-Goldner (EG) for evaluation of myelin loss, gliosis, and angiopathic changes, respectively. In addition, the sections were immunostained with antibodies against the following: glial fibrillary acidic protein (GFAP, polyclonal, 1:1,500, Dako); α-smooth muscle actin (SMA, monoclonal, 1A4, 1:50, Dako); phosphorylated neurofilament (pNF, monoclonal, SMI31, 1:1,000, Sternberger Monoclonals), an axon marker; collagen IV (COL4, polyclonal, 1:500, abcam), a marker of basement membrane within and adjacent to vessels. An antibody against the extracellular domain of Notch3 (Notch3 ECD, monoclonal, 1G5, 1:100, Abnova) was also used for the CADASIL group. Gallyas-Braak silver staining and antibodies against amyloid β 11-28 (monoclonal, 12B2, 1:50, IBL), phosphorylated tau (monoclonal, AT8,1:200, Fujirebio) and phosphorylated α-synuclein (monoclonal, pSyn#64, 1:1000, Wako) were used to assess senile pathologic changes based on the “ABC” score, CAA score and the fourth consensus report of the DLB consortium.^31–33^ Bound antibodies were visualized by the peroxidase-polymer-based method using a Histofine Simple Stain MAX-PO kit (Nichirei, Tokyo, Japan) with diaminobenzidine as the chromogen. Immunostained sections were counterstained with hematoxylin.

### Identification of the Regional Pathology in the WM

To quantify each pathological variable in an identical region, we divided the WM into 5-mm square areas (Fig 1B). The total numbers of divided areas in the CSO and temporal WM were 9.3±2.4 and 5.3±0.9 in the CADASIL group, 15.3±3.3 and 7.7±0.9 in the *HTRA1*-AD group, 10.7±1.7 and 5.0±0.8 in the sSVD group, and 17.7±2.1 and 7.7± 0.5 in the controls. Completely infarcted areas were excluded from the analysis because the patterns of alteration they contained were entirely different.^34^

### Assessment of White Matter Degeneration

For assessment of WM degeneration, we quantified and analyzed four variables in each patient: (1) myelin density, (2) axon density, and two pathologic changes related to astrocytes, i.e. (3) clasmatodendrocyte (CL) density and (4) extent of gliosis.

(1) For evaluation of myelin density, we used a modified form of the myelin index (mMyelin index).^34^ Using KB-stained sections, multiple images of each area were taken through a ×4 objective lens. The mean gray value for each divided area, corresponding to the staining intensity, was calculated using the NIH Image J software package (imagej.net). We then obtained the “mMyelin index” as a ratio relative to the value in a normal area in each subject (Fig 1B and C). (2) For evaluation of axon density, multiple images of each area in pNF-immunostained sections were taken through a ×10 objective lens. Using NIH Image J, the percentage of the pNF-immunoreactive area was measured within each field, and the value was averaged to obtain the percentage pNF area for all sections in each divided area. (3) For assessment of clasmatodendrocytes, we first evaluated GFAP-immunoreactive cells by dividing them into four types based on their morphology: apparently normal cells with long fine processes, hypertrophic reactive cells, cells that had swollen and rounded cytoplasm with beaded processes showing weak immunopositivity (clasmatodendrocytes),^14,15^ and fibrillary cells (gliosis). Because clasmatodendrocytes are often intermingled with gliosis, it was difficult to differentiate gliosis from clasmatodendrocytes only on the basis of GFAP immunostaining. Therefore, we used Holzer staining for specific visualization of gliosis as described below. To quantify CL density, we counted the numbers of GFAP-immunoreactive cells with a nucleus showing features of CL (Fig 1D to F), and expressed the value as density per unit area (CL number/mm^2^). (4) To determine the degree of gliosis, we employed quartile analysis (0; none, 1; mild, 2; moderate, 3; severe) of Holzer-stained sections (Fig 1G).

### Assessment of Vascular Degeneration

For assessment of vascular degeneration, we quantified and analyzed four variables related to vasculopathy in each patient: (1) SMA density, (2) arteriole sclerosis, (3) capillary sclerosis, and (4) extent of the perivascular space (PVS). In addition, Notch3 ECD density was assessed in the CADASIL group.

(1) To evaluate SMA and Notch3 ECD density, multiple images of each area in immunostained sections were taken through a ×4 objective lens. The percentage of the immunoreactive area was then calculated in a similar manner to the percentage area of pNF. (2,3) Arteriole and capillary sclerosis was determined as the sclerosis index (SI).^35^ Arterioles (50-200 μm in diameter) and capillaries (<10 μm in diameter) were observed using EG-stained sections under x10 and x40 objectives, respectively, and images of each area were taken. The outer and inner diameters of the vessels (Dext and Dint) were measured at two different points and the SI was calculated by incorporating the diameters into the devised formulae (Fig 1H and J). (4) The extent of the PVS was expressed as a PVS extent index (PVSI). To visualize the PVS clearly, COL4-immunostained sections were used and images of the PVS around arterioles were taken through a x10 objective. Then, the diameter of the PVS and the outer diameter of arterioles (Dpvs, Da) were measured at two different points and the PVSI was calculated in a similar manner to the SI (Fig 1I and J).

### Tissue Clearing-based 3D Analysis of Small Vessels and Astrocytic features

To clarify the spatial distribution of SMA loss within vessels and the spatial relationship between the abnormal features of astrocytes and vessels in the frontal CSO, we employed a 3D analytical method combined with a chemical tissue clearing technique. We prepared 10% formalin-fixed tissue blocks of approximately 1 cm^3^ of the frontal CSO (Fig 5A) from three patients with CADASIL (CAD 2 and 3 in Table 1, and a male aged 57 years, harboring Arg449Cys *NOTCH3* mutation), three patients with *HTRA1*-AD (*HTRA1*-AD 1-3 in Table 1), three patients with sSVD (sSVD 1-3 in Table 1), and nine controls (younger controls, Con 2, 4-7; and older controls, Con 8-11 in Table 2B). Because of the retrospective nature of the present study, the formalin fixation time varied from 1 to 20 years (mean: 6.5 years). The previously published clearing protocols (CUBIC^22^ and iDISCO^36^) were modified to satisfy both antigenicity and transparency for 3D imaging of human brain samples. Typically, each procedure involved 1) delipidation, 2) H_2_O_2_-based bleaching, 3) immunostaining, and 4) clearing (Fig 5B). Alexa Fluor 647-labeled anti-αSMA (monoclonal, 1A4 + ACTA2/791, 1:100, Novus Biologicals) and anti-laminin (polyclonal, 1:100, Novus Biologicals), and Cy3-labeled anti-GFAP (monoclonal, 1:100, Sigma Aldrich) antibodies were used for detecting vessels and astrocytes, respectively. The antibodies for laminin and GFAP were used for double-immunofluorescence labeling. We applied different delipidation cocktails for preserving the antigenicity of αSMA, laminin and GFAP. For αSMA-labeling, the brain blocks were immersed in 10% 1,2-hexanediol (Hxd)/1% sodium sulfite-PBS solution (pH 8.5) with shaking at 45°C for 3 days. For laminin and GFAP-labeling, the brain blocks were treated with 10% Hxd in distilled water with shaking at 45°C for 3 days. After washing in PBS, the brain blocks were bleached with 1% H_2_O_2_ in PBS with shaking at 45°C for 2 days. Each individual brain block was then placed in 1 ml of immunostaining buffer (mixture of PBS, 0.5% Triton X-100, 0.25% casein, and 0.01% NaN_3_) containing Alexa Fluor 647-labeled αSMA or Alexa Fluor 647-labeled laminin and Cy3-labeled GFAP for 1 weeks. After washing with PBS, the samples were cleared using CUBIC-R (an aqueous mixture of 45 wt% antipyrine and 30 wt% nicotinamide) or BABB (a 1:2 mixture of benzyl alcohol/benzyl benzoate) depending on the WM content of the sample. BABB was used for clearing of lipid-rich tissue samples. For CUBIC-R clearing, the samples were immersed in 1:1-diluted CUBIC-R with gentle shaking at room temperature overnight. The samples were then immersed in CUBIC-R with gentle shaking at room temperature for 1-2 days. For BABB clearing, the samples were dehydrated using steps of 60% (4 h) – 80% (4 h) – 100% (overnight) – 100% (4 h) methanol with gentle shaking at room temperature. The samples were then cleared using basic BABB containing 3% *N*-butyldiethanolamine with gentle shaking at room temperature overnight.

3D images of SMA-positive structures were captured using light-sheet fluorescence microscopy (LSFM: Olympus, MVX10-LS. SMA and laminin: ex637 nm, GFAP: ex532 nm, Autofluorescence: ex532 nm). Images were captured using a x0.63 objective lens [numerical aperture (NA) = 0.15, working distance = 87 mm] with a x1-6.3 digital zoom. When the stage was moved in an axial direction, the movement of the objective lens was also synchronized in an axial direction to avoid defocusing. The RI-matched sample was immersed in BABB or an oil mixture (RI = 1.525) composed of silicone oil HIVAC-F4 (RI = 1.555, Shin-Etsu Chemical Co., Ltd.) and mineral oil (RI = 1.467, M8410, Sigma-Aldrich) for CUBIC-R clearing during image acquisition.

### Genetic Analysis

Genomic DNA was extracted from the patients’ peripheral lymphocytes or fresh-frozen frontal samples. We used the primer pairs for the *HTRA1* and *NOTCH3* genes that have been described previously.^18^

### Statistical Analysis

Mean and Standard deviation (SD) were used to describe the distributions of continuous variables. The data for the frontal CSO and temporal lobe were analyzed separately. Analysis of variance was performed to compare the means of pathological variables among the four groups. The Tukey-Kramer procedure was used as a post hoc test for correction of the significance level for avoiding any excess of false positivity. Spearman correlation analysis was performed to evaluate correlations between the variables studied. Data were analyzed using the JMP version 15.0 software package (SAS Institute, Cary, NC, USA). The level of statistical significance was set at *P* <0.05.

## Results

### White Matter Degeneration

#### Myelin and Axon Densities

KB-stained sections revealed loss of myelinated fibers and depletion of oligodendrocytes, which varied across the groups (Fig 3A). The mMyelin index based on examination of KB-stained sections increased with the severity of WM pathology. As expected, the scores for the controls (CSO: 1.08±0.039, temporal: 1.00±0.078) were significantly lower than those for the CADASIL (1.57±0.22, p <0.0001; 1.54±0.22, p <0.0001), *HTRA1*-AD (1.14±0.036, p = 0.014; 1.13±0.037, p = 0.19) and sSVD (1.20 ±0.099, p <0.0001; 1.17±0.024, p = 0.044) groups, except for the temporal WM in *HTRA1*-AD. In both the CSO and temporal WM, scores were significantly maximal in the CADASIL group, followed in order by sSVD (CADASIL vs sSVD, CSO: p <0.0001 and temporal: p = 0.00030) and then *HTRA1*-AD (CADASIL vs *HTRA1*-AD, p <0.0001 and p <0.0001) (Fig 3A’).

**Figure 3:**
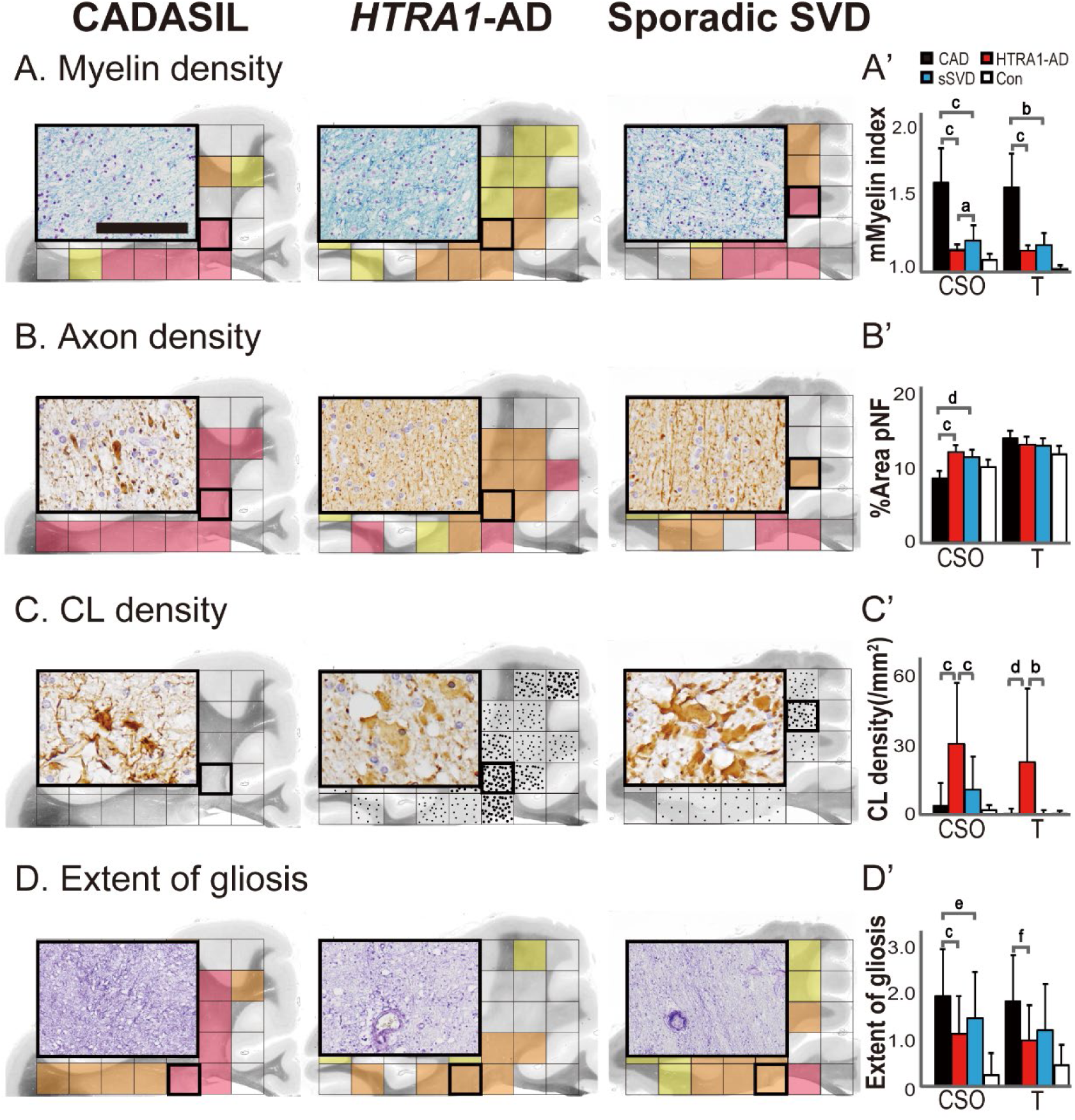
Quantitative analyses of white matter degeneration. **(A-D)** Histopathology of the white matter in the frontal centrum semiovale (CSO) in representative patients with CADASIL (CAD 2 in Table 1), *HTRA1*-AD (*HTRA1*-AD 1) and sporadic SVD (sSVD 3). Schematic representation of the distribution of degeneration from a single case based on quartilized data for each unit area. Each dot represents the approximate number of clasmatodendrocytes **(C)**. White square, none (first quartile - 25^th^ percentile); yellow square, mild (median value - 50^th^ percentile); orange square, moderate (third quartile - 75^th^ percentile); red square, severe (>75^th^ percentile). **(A’-D’)** Mean (±SD) value of the modified Myelin index, %area pNF, numbers of clasmatodendrocytes and the extent of gliosis in the frontal CSO and temporal white matter. CSO, central semiovale; T, temporal white matter; a, p = 0.014; b, p = 0.00030; c, p <0.00010; d, p = 0.0012; e, p = 0.029; f, p = 0.0060. Bar = 150 μm **(A)**, 60 μm **(B, C)** and 400 μm **(D)**.

pNF immunohistochemistry revealed features of severe axonal degeneration, such as abundant swollen and fragmented axons showing an irregular arrangement in the CSO of patients with CADASIL (Fig 3B, *left panel*), whereas axons were relatively preserved, even in the CSO, in patients with *HTRA1*-AD and sSVD, similar to the situation seen in controls (Fig 3B, *middle and right panels*). Indeed, the pNF-immunoreactive areas in the CSO in CADASIL were significantly decreased in comparison to *HTRA1*-AD and sSVD (8.31±3.81 vs 11.73±3.78, p <0.0001 and 11.13 ±2.62, p = 0.0012), whereas no significant difference was evident in the temporal lobes (13.61±2.33 vs 12.76±2.66, p = 0.66 and 12.62±1.65, p = 0.55) (Fig 3B’).

#### Astrocytic Features: Clasmatodendrocytes and Gliosis

GFAP immunohistochemistry demonstrated various changes in astrocytes among the groups. Surprisingly, there was a noticeable increase of CL density [number/mm^2^] in the deeper WM in both the frontal CSO and temporal WM in the patients with *HTRA1*-AD (Fig 3C, *middle panel*), as compared with those in the CADASIL and sSVD groups (CSO: 29.21±25.64 vs 3.92±9.43, p <0.0001, and 10.22±14.12, p <0.0001, temporal: 21.26±31.13 vs 0.63±2.00, p = 0.0012 and 0.36±1.71, p = 0.00030) (Fig 3C). Indeed, no CL was detected in 2 of the 3 patients with CADASIL who had a longer disease duration, although severe gliosis was evident (Fig 3C, *left panel*). In the patients exhibiting moderate gliosis with an increase in CL number, most of whom were in the sSVD group, the CL processes appeared to have changed from a beaded to a thicker fibrillary form, and the remaining cytoplasmic body was swollen (Fig 3C, *right panel*). The corresponding CL densities in the CSO and temporal WM did not differ significantly among the CADASIL, sSVD and control groups (CADASIL vs controls and sSVD vs controls, CSO: p = 0.34 and p = 0.053, temporal WM: p = 1.0 and p = 1.0) (Fig 3C’).

The distribution of gliosis visualized by Holzer staining was similar in the CADASIL, *HTRA1*-AD and sSVD groups, being fibrillary and diffuse with dark purple staining, prominently around the arterioles in the periventricular and deep WM (Fig 3D), resembling the distribution of leukoaraiosis evident on T2-weighted MRI. The extent of gliosis varied in relation to the disease state of each patient in each group, but quartile analysis revealed that it was most severe in CADASIL irrespective of the disease state, followed in order by sSVD, and then *HTRA1*-AD in both the CSO and temporal WM (CSO: 1.9±0.8 vs 1.4±0.8, p = 0.029 and 1.1±0.8, p <0.0001, temporal WM: 1.8±1.0 vs 1.2±0.8, p = 0.073 and 1.0±0.8, p = 0.0060) (Fig 3D’).

### Vascular Degeneration

#### SMA density

Smooth muscle cells in the tunica media within the arteriole and wall cells (i.e. pericytes) surrounding the capillaries showed positive immunoreactivity for SMA. There appeared to be a markedly reduced area of SMA immunopositivity in the patients with CADASIL and *HTRA1*-AD, whereas in those with sSVD the area was retained to a relative degree (Fig 4A). We also observed diffuse loss of SMA within the vessels distributed in the affected WM, involving both capillaries and arterioles in CADASIL, and severe loss in arterioles with relative preservation in capillaries in *HTRA1*-AD (Fig 4A, *middle panel*, *arrow*). Indeed, the percentage area of SMA staining in the frontal CSO was significantly lower in *HTRA1*-AD than in the CADASIL and sSVD groups (0.24±0.072 vs 0.42±0.16%, p=0.026, and 0.60±0.38, p<0.0001), and in the temporal WM in the sSVD group (0.26±0.11 vs 0.43±0.17, p=0.19, and 0.53±0.16, p=0.0032) (Fig 4A’), although the present quantitative analysis revealed milder degeneration of the WM in the *HTRA1*-AD group than in the CADASIL and sSVD groups.

**Figure 4:**
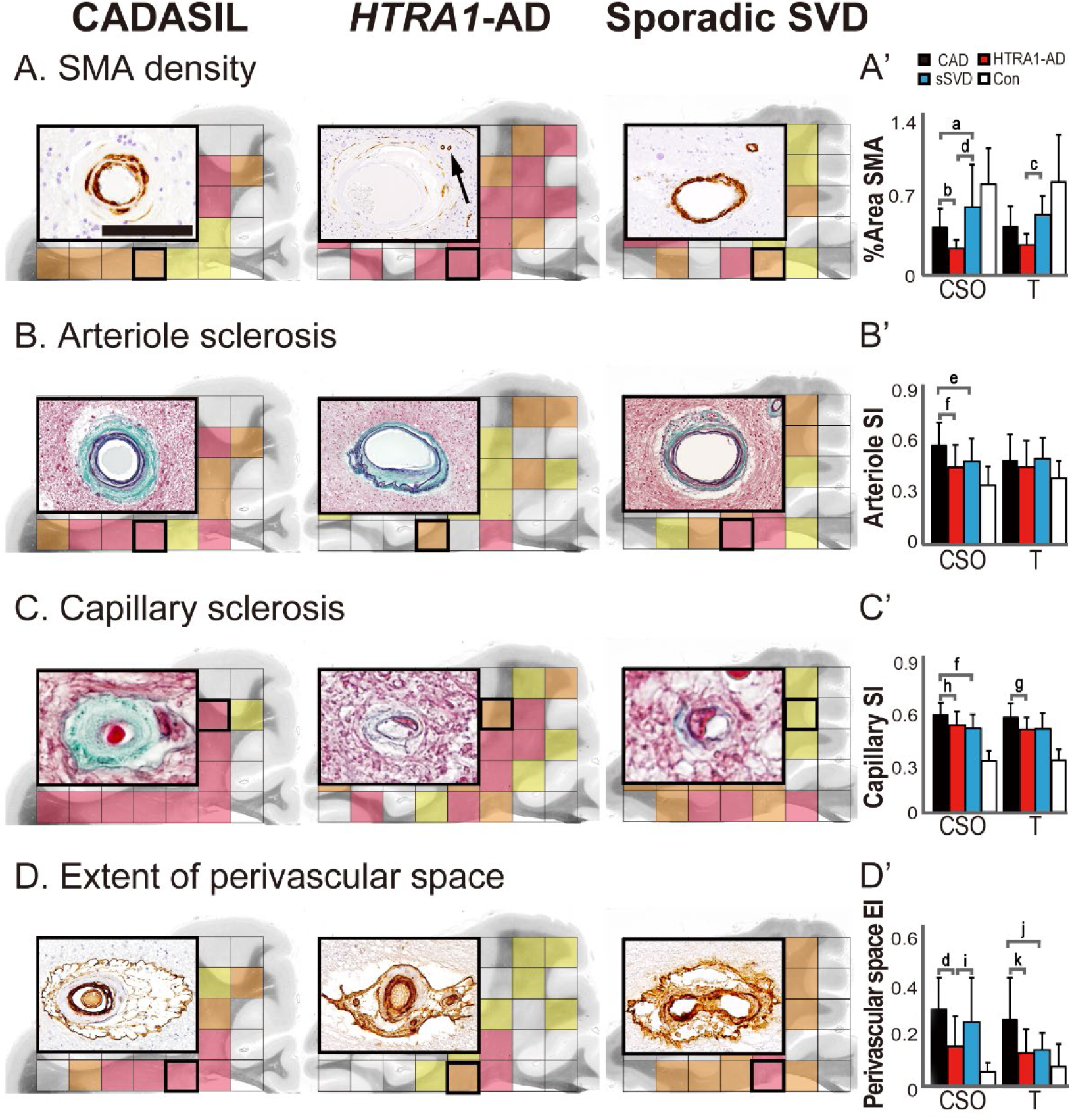
Quantitative analyses of the vascular degeneration. **(A-D)** Histopathology of vasculopathy in the frontal centrum semiovale (CSO) in representative patients with CADASIL (A, B, D; CAD 3 and C; CAD 2 in Table 1), *HTRA1*-AD (A; *HTRA1*-AD 3 and B-D; *HTRA1*-AD 2) and sporadic SVD (A-C; sSVD 4 and D; sSVD 2). Schematic representation of the distribution of degeneration from a single case based on quartilized data for each unit area. White square, none (first quartile - 25^th^ percentile); yellow square, mild (median value - 50^th^ percentile); orange square, moderate (third quartile - 75^th^ percentile); red square, severe (>75^th^ percentile). **(A’-D’)** Mean (±SD) value of %area SMA, arteriole and capillary sclerosis index, and PVS extension index in the frontal CSO and temporal white matter. CSO, central semiovale; T, temporal white matter; SI, sclerosis index; EI, extension index; a, p = 0.040; b, p = 0.026; c, p = 0.0032; d, p <0.0001; e, p = 0.019; f, p = 0.00020; g, p = 0.048; h, p = 0.0047; i, p = 0.0013; j, p = 0.0012; k, p = 0.00040. Bar = 20 μm **(A)**, 200 μm **(B)**, 15 μm **(C)** and 300 μm **(D)**.

#### Sclerosis Index

We examined cross-sections of approximately 520 and 770 consecutively selected vessels to calculate the SI of arterioles and capillaries in the frontal CSO and temporal WM. The SI was well consistent with the degree of collagenous wall thickening assessed using HE and EG staining (Fig 4B, C). In both the hereditary and sporadic SVD groups, vessels had a higher SI than those in the controls. The SI of arterioles and capillaries in the frontal CSO was significantly higher in CADASIL than in sSVD and *HTRA1*-AD (arteriole SI: 0.56±0.13 vs 0.48±0.13, p=0.019 and 0.44±0.12, p=0.0002, capillary SI: 0.57±0.072 vs 0.50±0.083, p=0.0002 and 0.51±0.081, p=0.0047) (Fig 4B’ and C’). In the temporal WM, the arterioles also appeared to be most severely affected in CADASIL, but differences among the disease groups did not reach statistical significance (p=0.99 and p=0.30) (Fig 4B’). In contrast, there was a significant difference in capillary SI between the CADASIL and *HTRA1*-AD groups, even in the temporal WM (0.56±0.087 vs 0.49±0.069, p=0.048) (Fig 4C’).

#### PVS Extension Index

The perivascular space is the space between penetrating vessels and the pia mater, where COL4 immunostaining clearly visualized the outer boundary of the pia mater (Fig 4D). We examined cross-sections of approximately 520 consecutively selected arterioles to calculate the PVS extension index. In contrast to the punctate or linear structures evident on MRI, considered to be enlarged PVS, in *HTRA1*-AD and sSVD, the apparent dilation was evident in CADASIL (Fig 4D, *left panel*) in both the frontal CSO and temporal WM, where the value of the PVS extension index in CADASIL was significantly higher than in *HTRA1*-AD (CSO: 0.31±0.13 vs 0.16±0.12, p<0.0001, temporal: 0.27±0.16 vs 0.14±0.094, p=0.0004). The index in the temporal WM also differed significantly between CADASIL and sSVD (0.15±0.071, p=0.0012), but did not reach significance in the frontal CSO (p=0.34) (Fig 4D’).

#### Tissue Clearing-based 3D Analysis

Our clearing/immunostaining protocol involved 1) delipidation (including permeabilization and antigen retrieval)/bleaching (including suppression of autofluorescence), 2) immunostaining, and 3) optically clearing. The potential sensitivity and specificity of immunolabeling are mainly dependent on the degree of delipidation. Post-fixation and clearing may enhance or suppress non-specific signals. In CUBIC-R clearing, the final transparency of the tissue depends on the stringency of delipidation, but MeOH dehydration/BABB mounting achieves efficient clearing of myelin-rich tissues that retain abundant lipid even after mild delipidation. On this basis, we established a series of 3D imaging protocols for use in this study.

#### SMA Immunohistochemistry

Immunohistochemistry for SMA clearly detected vessels >20 μm in diameter, the signals being more intense in arteries than in veins, and successfully demonstrated them as 3D structures (Fig 5C). Within 1-cm^3^ blocks of the frontal CSO, 5-10 main medullary arteries (100-300 μm in diameter) ran parallel in the direction of nerve fibers, each separated by about 1 mm, forming several branches and together constituting a vascular net (Fig 5C; *inset*).

Interestingly, in the disease groups, two distinct patterns of SMA loss were visualized within the vessels distributed in WM lesions: (1) CADASIL and sSVD: diffuse loss, being particularly prominent in small branches; (2) *HTRA1*-AD: selective loss in main branches. In CADASIL and sSVD, segmental loss of SMA within arteries showing tortuous courses was diffusely evident in the frontal CSO (Fig 5D and E, *arrowhead*), and this was accentuated in smaller branches (<50 μm), leading to a depletion of SMA with sparse preservation of SMA in the main branches in the patient with advanced CADASIL (Fig 5D). Moreover, in CADASIL, the diameter of the vessels appeared to be considerably greater, due to the thickened intima, forming its inner component. In contrast, main branches were predominantly affected in *HTRA1*-AD (Fig 5F, *arrowhead*), while SMA immunopositivity became evident in contiguous distal areas (Fig 5G, *arrow*).

**Figure 5:**
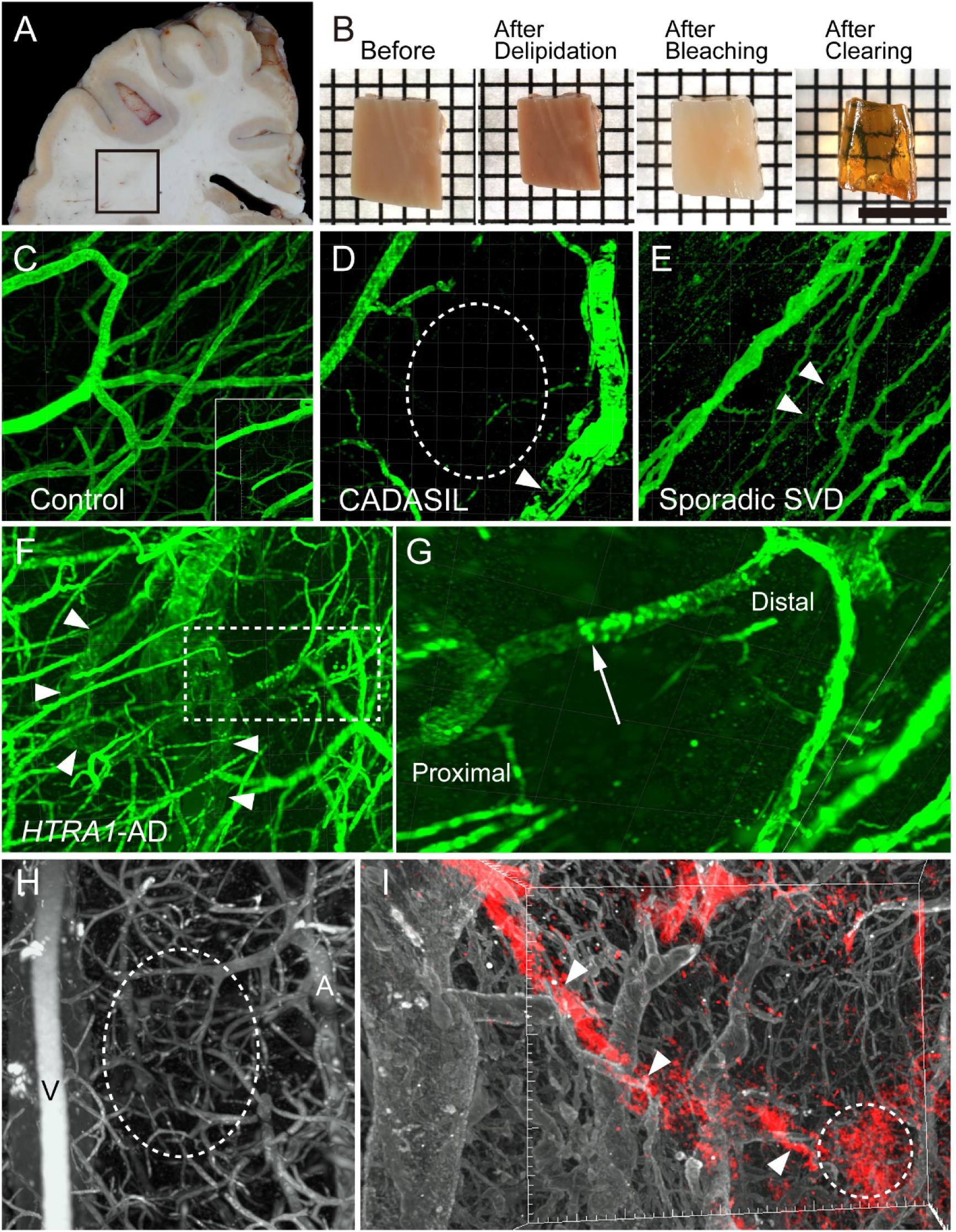
Appearance of a brain block under chemical tissue clearing and 3D images of vascular networks in representative patients. **(A)** The representative location of a block used for 3D analysis in the frontal centrum semiovale (CSO) of a patient (sSVD1 in Table 1). Formalin-fixed brain. **(B)** Appearance of a 1-cm^3^ brain block before (in PBS) and after each chemical clearing process. After Delipidation, delipidation by 10% HxdHx at 45°C for 3 days; After Bleaching, bleaching by 1% H_2_O_2_ at 45°C for 2 days; After Clearing, clearing by dehydration followed by BABB. **(C-G)** A coronal view of 3D vascular networks demonstrated by Alexa Fluor 647-labeled anti-SMA antibody. Frontal CSO. Abundant small arterioles of a control (Con 7 in Table 2B) in coronal view appear to be running parallel in sagittal view (*inset*) **(C).** SMA loss is more accentuated within the smaller branches in a peripheral area (*circle*) in CADASIL (CAD 2 in Table 1) **(D)**, and sporadic SVD (sSVD 1 in Table 1). Spiral twisting of the vessels is also evident **(E)**. In contrast, segmental SMA loss is demonstrated within predominantly main branches (*arrowheads*) in a patient with *HTRA1*-AD. (*HTRA1*-AD 3 in Table 1) **(F)**. The magnified image shows the retained SMA in the distal part (*arrow*) of same branch **(G)**. **(H, I)** 3D pathology demonstrated by double immunofluorescence staining for laminin and GFAP. Abundant small vessels visualized by Alexa Fluor 647-labeled anti-laminin antibody including arterioles, venules and capillaries in a control (Con 7 in Table 2B) **(H)**. Gliosis (red) along the arterioles (*arrowhead*) and within a peripheral area (*circle*) **(I)**. V, veins; A, arterioles. Bar = 1.5 cm **(A)**, 7 mm **(B)**, 1 mm **(C-F),** 150 μm **(G),** 400 μm **(H)** and 700 μm **(I)**.

#### Laminin and GFAP Immunofluorescence

Immunofluorescence staining for laminin clearly detected vessels >5 μm in diameter, and equally visualized both arteries and veins, which were more numerous than the small vessels demonstrated by SMA immunofluorescence. The spaces among main branches (peripheral area) were occupied by capillary loops (Fig 5H, *circle*). On the other hand, immunofluorescence staining for GFAP demonstrated normal astrocytes and gliosis, although the signal intensity of clasmatodendrocytes was too weak to be visible. Therefore, we applied double-labeling immunofluorescence for GFAP and laminin to observe the spatial positional relationship between gliosis and vessels including capillaries. This revealed fuzzy gliosis within the peripheral areas (Fig 5I, *circle*) in addition to that abundantly evident around the arterioles (Fig 5I, *arrowheads*), the latter being consistent with the results of Holzer staining (Fig 3D). This phenomenon was most apparent in the patients with CADASIL.

## Discussion

In the present study, we quantitatively demonstrated that CADASIL showed the most aggressive pathology, with unique dynamics of clasmatodendrosis and SMA loss in *HTRA1-*AD, among sporadic cases and two representative SVDs. Furthermore, we developed a 3D imaging method for visualization of SMA and laminin-labeled vasculature even in the cerebral WM, and found two distinct patterns of SMA loss in the WM: (1) diffuse SMA loss apparent in smaller branches in CADASIL and sSVD; (2) selective involvement of the main branches in *HTRA1*-AD.

Consistent with the present findings, several histopathological studies of the WM of patients with SVDs have demonstrated that CADASIL has the most aggressive pathology such as more profound myelin loss and a higher density of degenerated axons.^35,37^ It also has the most severe degree of arteriolar sclerosis, a decreased area of SMA immunoreactivity, and extension of the PVS relative to sporadic and other hereditary SVDs including pontine autosomal dominant microangiopathy and leukoencephalopathy, and hereditary endotheliopathy with retinopathy, nephropathy, and stroke.^7^ However, the present study has revealed for the first time that in comparison with CADASIL, *HTRA1*-AD has a considerably higher number of clasmatodendrocytes and more profound SMA loss, despite the fact that the WM and small vessels are less severely affected.

Clasmatodendrosis in the WM has been described in various situations,^14,15,38–40^ particularly brain edema,^38–40^ where some markers of autophagy such as p62 and LC3^14,15,38,39^ have been revealed in autopsied brain tissue, suggesting an irreversible autophagic phase. One study using a non-human primate model demonstrated that cerebral hypoperfusion caused by 3-vessel occlusion led to increased fibrinogen immunoreactivity suggestive of capillary leakage, and an increase in the number of clasmatodendrocytes until 14 days after occlusion, followed by a decline of both parameters as the expression of capillary markers increased.^14^ These findings indicate that clasmatodendrocytes emerge transiently in the early phase in response to hypoperfusion due to disruption of the gliovascular unit, and are then disrupted by autophagy. Indeed, in the present study, moderate numbers of clasmatodendrocytes were observed in the frontal CSO and temporal WM in CADASIL 1, but not in CADASIL 2 and 3 where the disease duration had been longer, and an extensive compensatory glial response (gliosis) had occurred instead.

In contrast, regardless of disease stage, the number of clasmatodendrocytes in each patient with *HTRA1*-AD was considerably higher than that in the patients with other SVDs. As HTRA1 is highly expressed in astrocytes as well as vasculature in the human brain, it can be speculated that these changes in the characteristics of astrocytes caused by mutation of *HTRA1* are attributable to such profound clasmatodendrosis. In fact, it has been reported that loss of HtrA1 in cultured astrocytes from mouse brain alters the nature of astrocytes, including their differentiation and response to injury.^41^ However, the association between astrocytes harboring mutation and clasmatodendrosis has remained unclear. With regard to the factors associated with clasmatodendrosis in patients with *HTRA1*-AD, we found that the percentage area of SMA staining was closely and positively correlated with the density of clasmatodendrocytes (CSO: ρ = 0.47, p = 0.0009, temporal: ρ = 0.63, p = 0.0014).

Proteomic analysis of cerebral vessels in human patients and model mice has suggested that CADASIL and CARASIL share a similar pathomechanism involving functional loss of the protease HTRA1.^42^ However, the present analysis revealed a distinct difference in the distribution of SMA loss within the vessels between the two diseases, sporadic SVD apparently showing a CADASIL-type pattern. With regard to SMA loss, the results of our integrated 2D and 3D analysis suggested that the markedly lower percentage area in *HTRA1*-AD reflects selective involvement of the main trunks of arterioles, which accounts for a large proportion of SMA immunoreactivity, from an early disease stage. In contrast, in the autopsied brain of CADASIL patients and also in model mice, the Notch3 ECD has been shown to accumulate preferentially in capillaries and a reduction of capillary density has been demonstrated.^43,44^ This is consistent with our finding that capillaries were markedly affected in CADASIL, accompanied by gliosis in the peripheral area of the vascular net, which is composed mainly of capillaries, while being relatively well preserved in *HTRA1*-AD. Similarly, although more mildly than in CADASIL, sporadic SVD showed a degenerative pattern involving both the main trunk and capillaries, i.e. the “smaller branches type”. Although the reason for the vulnerability of larger arterioles in *HTRA1*-AD is unclear, larger-size cerebral arteries are reportedly affected in patients with CARASIL^19,20^ and in visceral organs model mice.^45^ As vascular smooth muscle cells of arteriolar and pre-capillary arterioles maintain a contractile tone and control the regulation of blood pressure and redistribution of blood flow,^46^ disruption of SMC within the main trunk may induce extensive capillary leakage and edema, which could affect the gliovascular unit and increase the number of clasmatodendrocytes. Further studies are needed to clarify the significance of this specific vulnerability of larger arterioles and its relationship to clasmatodendrosis in the context of *HTRA1*-related gliovascular degeneration.

Considering the process of WM degeneration in relation to vascular changes, the present quantitative analysis revealed that the mMyelin index was a highly sensitive marker of temporal WM change in the CADASIL group, even from an early disease stage. Thus, to clarify the essential aspects of vasculopathy affecting early myelin loss, we investigated several variables within the temporal WM, but found that none were significantly correlated with the mMyelin index, including the percentage area of Notch3 ECD in the CADASIL group. Similarly, no such variable was evident in the sSVD and *HTRA1*-AD groups, except for a positive correlation with the degree of PVS extension in the sSVD group (ρ = 0.44, p = 0.042). On the other hand, in the frontal CSO, where lesions were more advanced, myelin density was correlated with the arteriole sclerosis index in the CADASIL group (ρ = 0.42, p = 0.029), and with the arteriole sclerosis index and PVS extension index in the *HTRA1*-AD group (ρ = 0.44, p = 0.0025 and ρ = 0.34, p = 0.0020). There were no significantly correlated vasculopathy variables in the sSVD group. These findings are consistent with an early role of PVS dilation and potential exacerbation of arteriolar sclerosis, as has been reported previously.^47^ Based on this interpretation, the perivascular edema evident in earlier stages due to failure of interstitial fluid drainage would damage perivascular tissue and astrocytes, resulting in myelin loss and clasmatodendrosis, consistent with some of the pathological features in the WMH revealed by imaging.^2,14,39,40^ At the advanced stage, loss of autoregulatory contractility due to vascular remodeling such as alterations of stiffness within arterioles would lead to further tissue rarefaction, lumen occlusion and eventually ischemic necrotic foci followed by gliosis. Moreover, as observed in CADASIL, preferential involvement of capillaries would damage the surrounding tissue more directly and severely, in view of its crucial role in BBB function in the termini of arteriolar networks. This difference in capillary burden could be one reason for the variations in the severity of WM pathology evident among CADASIL, *HTRA1*-AD and sSVD, highlighting again the key role of capillaries and the BBB in disease progression.

One limitation of the present study was its small sample size. In addition, the present series of patients did not include any with CARASIL and *HTRA1*-AD with other mutations of *HTRA1*. Therefore, it may be important to confirm whether selective involvement of larger arterioles is evident in patients with CARASIL, and *HTRA1*-AD harboring other mutations.

## Conclusions

The present study has provided evidence that small vessels are principally affected by the pathological process associated with WM degeneration in the frontal centrum semiovale and temporal WM in patients with CADASIL, *HTRA1*-AD and sporadic SVD. Moreover, 3D immunohistochemistry revealed that SMA within the arteries of the WM have diffuse involvement including smaller branches in CADASIL and sporadic SVD, with more selective involvement of the larger main branches in *HTRA1*-AD, suggesting an association with the various distinct forms of disease progression shown by these SVDs. Further studies are needed to clarify the pathomechanism responsible for the distinct distribution of cerebral small vessel involvement underlying sporadic and hereditary SVDs in the context of disease development. The 3D analytical method we have developed may contribute to high-throughput analysis with cellular-level resolution for spatial assessment of neurogliovascular relationships in the human brain.

## Acknowledgements

This work was supported by Grant-in-Aid for Research Activity Start-up (JSPS KAKENHI Grant Number JP19K21314 to R.S.); Grant-in-Aid for Early-Career Scientists (JSPS KAKENHI Grant Number JP20K16595 to R.S.); Grant-in-Aid for Scientific Research(A) (JSPS KAKENHI Grant Number JP19H01061 to A.K.); Grant-in-Aid for Scientific Research(B) (JSPS KAKENHI Grant Number JP18H02105 to K.T.); Grant-in-Aid for Challenging Research (Pioneering, JSPS KAKENHI Grant Number JP20K20468 to A.K.) and (Exploratory, JSPS KAKENHI Grant Number JP18K19373 and JP20K21246 to K.T.); Grant-in-Aid for Scientific Research on Innovative Areas (MEXT/JSPS KAKENHI Grant Number JP20H04700 to K.T.); Grant-in-Aid from the Cell Science Research Foundation (K.T.); AMED Grant Number JP21zf0127004, JP21ek0210125, and JP21wm0425001 to K.T. The authors thank Yuko Sakaki (Department of System Pathology for Neurological Disorders, Brain Research Institute, Niigata University) for the expert technical assistance.

## Authors’ Contributions

R.S., K.T., and A.K. contributed to the conception and design of the study. All authors contributed to the acquisition and analysis of data. R.S., K.T., and A.K. contributed to the drafting of the manuscript and preparation of figures and tables.

